# Isocaloric restriction of a high-fat diet under TAK-242 administration: metabolic and gut microbial responses in mice

**DOI:** 10.64898/2026.11.05.764622

**Authors:** Eunji Jeong, Seoyeon Chae, Sungjin Eun, Hyunwoo Chae, Sunhye Lee

## Abstract

High-fat feeding promotes obesity in mice by increasing energy intake, so diet composition and energy intake are hard to separate without pair-feeding. We previously showed that isocaloric restriction normalizes body weight, glycemia and plasma lipids but not visceral adiposity, adipose inflammation or the fecal microbial community. We hypothesized that, during administration of the TLR4 signaling inhibitor TAK-242, matching energy intake to that of low-fat controls would normalize adiposity, intestinal barrier gene expression and the fecal microbial community. Male C57BL/6 mice (n = 8 per group) received a low-fat diet, a high-fat diet *ad libitum*, or a high-fat diet pair-fed to controls from week 0 or week 5 for 12 weeks; TAK-242 (3 µg/g, three times weekly) was given to every group from week 0.4, with no vehicle-injected arm. Intake did not differ among groups during a three-week low-fat acclimation (P = .88); over the 12 weeks on the assigned diets the *ad libitum* high-fat group ate 5.5% more than controls (P = .50). Over the full trajectory the *ad libitum* high-fat group gained 0.48 g/week more than controls (group × week, P = .012); both pair-fed groups had lower fasting glucose than that group. Adipocyte diameter, visceral fat proportion, ileal barrier genes, CCK-induced satiation and microbial alpha diversity did not differ significantly among groups. Adipose *Ccl2* expression differed (P = .043, exploratory), as did fecal community composition (PERMANOVA P = .022), although no pairwise contrast survived correction. Energy restriction did not return the microbial community toward the low-fat profile.

## 1. Introduction

Dietary fat promotes obesity in rodents chiefly by increasing how much animals eat rather than by changing how efficiently they convert food to tissue [1]. This makes the standard high-fat-diet model ambiguous: mice fed such a diet *ad libitum* differ from controls both in the composition of what they eat and in the energy they consume, and the two cannot be separated by observation. The gut microbiota is itself a regulator of feeding behavior [2], which makes the distinction material. Yoked pair-feeding resolves the ambiguity by holding energy intake equal while composition differs, and it can be imposed from the start of feeding, as a prevention model, or after obesity is established, as a treatment model.

We previously applied this design [3] and found that the two settings differ in what energy restriction can undo. Body weight, glycemia and plasma lipids followed energy intake and were normalized by restriction. The proportion of body weight stored as visceral fat, adipose *Tnf* expression in mice with established obesity, and fecal microbial diversity did not: they remained associated with diet composition despite matched energy. A normal body weight on a high-fat diet therefore did not imply normal adiposity, and the microbial community did not return toward the low-fat profile. Similar persistence of obesity-associated adipose phenotypes after weight loss has been reported in immune, transcriptional and epigenetic terms [4–6], and persistent microbial alterations accelerate weight regain after dieting [7].

One candidate link between dietary fat and the adipose phenotype that restriction fails to reverse is Toll-like receptor 4. Saturated fatty acids can activate TLR4 signaling independently of obesity [8], and lipopolysaccharide absorbed during high-fat feeding has been proposed to drive adipose inflammation through the same receptor [9, 10]. TLR4 signaling may therefore contribute to the persistence of adipose and microbial alterations under high-fat feeding, although its role when energy intake is restricted remains incompletely understood.

Here we repeated the prevention and treatment design while administering the small-molecule TLR4 signaling inhibitor TAK-242 [11, 12] to every animal throughout the study. Two questions motivated the repetition. First, each of the features that resisted energy restriction in our earlier work was observed in a single cohort, and replication in an independent cohort is a precondition for interpreting them. Second, it was unknown whether those features persist when TLR4 signaling is pharmacologically inhibited in every animal, a condition under which the contribution of diet composition can be examined against a common pharmacological background. Because the inhibitor was given to all four groups and no vehicle-injected arm was included, this experiment characterizes the dietary contrast during continuous TAK-242 administration rather than testing the contribution of TLR4 itself; we state this limitation at the outset and return to it in the Discussion. We asked which of the features that energy restriction failed to normalize previously [3] — adiposity, adipose inflammatory gene expression, intestinal barrier gene expression, CCK-induced satiation and the fecal microbial community — differed between dietary groups under that condition.

## 2. Methods and materials

### 2.1. Animals and ethics

Thirty-two male C57BL/6 mice were obtained from Samtako (Osan, Republic of Korea) at six weeks of age on 30 June 2023. Each animal was caged singly for the duration of the study in a room held at 23 ± 1 °C and 50 ± 5% relative humidity on a 12-h light/dark schedule, with water available at all times. Mice remained on the low-fat diet for three weeks before group assignment on 20 July 2023, which defines week 0 of the study. The protocol was approved by the Institutional Animal Care and Use Committee of Sun Moon University (approval no. SM-2023-01-01, approved 14 July 2023), and this report follows the ARRIVE guidelines [13].

### 2.2. Diets

Two purified diets were used, matched at 20% of energy from protein and differing in fat (10% versus 45% of energy), following D12450J and D12451 formulations (Supplementary Table S3). Calculated energy densities were 3.85 kcal/g for the low-fat diet and 4.73 kcal/g for the high-fat diet; these values were used to convert all intake measurements to energy, with the low-fat value applied to every group during the three acclimation weeks, when all mice ate the low-fat diet. Both diets were the same preparations used in our preceding experiment [3].

### 2.3. Experimental design

Allocation used body-weight strata: the 32 animals were ranked, divided into strata, and assigned at random within each stratum to one of four groups of eight on 20 July 2023, which defines week 0 of the study (Fig. 1A). The assigned diets were provided from 21 July 2023 and the animals were followed for 12 weeks, to 13 October 2023. The LF group stayed on the low-fat diet. The HF group ate the high-fat diet without restriction for the whole period. The HF/PF group ate the high-fat diet without restriction for five weeks and was then restricted from week 5 (25 August 2023), modeling treatment of established obesity. The PF/PF group was restricted from 22 July 2023, the day after the assigned diets began, modeling prevention. Restriction of the PF/PF group therefore began two days before the first TAK-242 injection (24 July 2023), and the design cannot separate the two. Weighing was weekly; food was weighed daily and converted to energy with the densities given above. No animal died or was removed from analysis.

**Figure 1.**
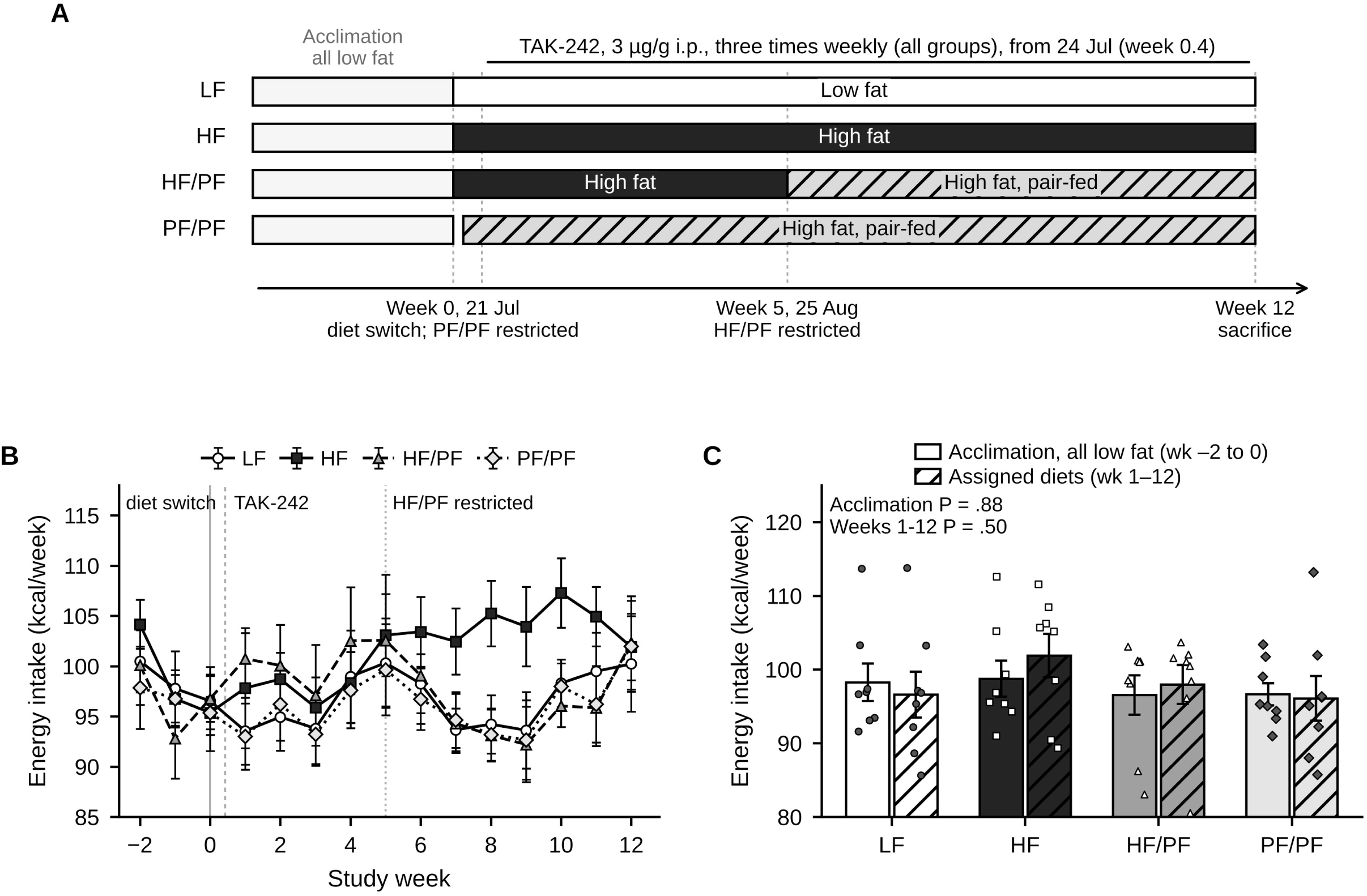
Study design and energy intake. (A) Design as performed. All four groups received TAK-242 (3 µg/g body weight) by intraperitoneal injection three times weekly from 24 July 2023 (week 0.4) until sacrifice; no vehicle-injected arm was included. LF received a low-fat diet throughout. HF received a high-fat diet *ad libitum* throughout. HF/PF received a high-fat diet *ad libitum* for five weeks and was then pair-fed to the low-fat group from week 5 (treatment model). PF/PF was pair-fed from 22 July 2023, the day after the diets were assigned (prevention model). Weeks −2 to 0 are the acclimation period, during which every mouse ate the low-fat diet; intake over those weeks was converted to energy with the low-fat density. (B) Weekly energy intake per mouse. Symbols are group means ± SEM; the solid line marks the diet switch at week 0, the dashed line the start of TAK-242 injections and the dotted line the start of restriction in HF/PF at week 5. (C) Energy intake averaged over the acclimation weeks (−2 to 0) and over the 12 weeks on the assigned diets (weeks 1–12). Bars are mean ± SEM with individual mice overlaid; P values are one-way ANOVA. Throughout the figures: LF, open bars and circles; HF, filled black bars and squares; HF/PF, dark grey bars and triangles; PF/PF, light grey bars and diamonds. n = 8 per group unless stated otherwise.

### 2.4. TAK-242 administration

All four groups received the TLR4 signaling inhibitor TAK-242 (resatorvid; 3 µg/g body weight, equivalent to 3 mg/kg; cat. no. HY-11109, MedChemExpress, Monmouth Junction, NJ, USA) by intraperitoneal injection three times weekly (Monday, Wednesday and Friday) until sacrifice. The dose follows the only published in vivo regimen for this compound in male C57BL/6 mice, in which 3 mg/kg was given intraperitoneally [14]; that study used a single injection in an endotoxemia model, so the repeated thrice-weekly schedule over 12 weeks used here has no direct precedent. Target engagement was not verified: neither downstream TLR4 signaling nor plasma TAK-242 concentrations were measured, so the degree to which TLR4 signaling was inhibited, and whether any inhibition persisted across 12 weeks of thrice-weekly dosing, is unknown. We therefore describe the condition throughout as continuous TAK-242 administration, and nowhere as sustained TLR4 blockade. A vehicle-injected arm was not included: the inhibitor was adopted in place of a genetic *Tlr4* knockout for reasons of availability and cost, and the available animals were allocated to the four dietary groups. Consequently this study cannot isolate the contribution of TLR4 inhibition, and all comparisons reported here are between dietary groups during continuous TAK-242 administration. The comparison with the preceding experiment [3], in which no inhibitor was given, is discussed as an observation across experiments rather than as a controlled contrast.

TAK-242 was prepared as a 2 mg/mL stock by dissolving the vial contents in dimethyl sulfoxide and making the volume up with physiological saline to give a final vehicle of 4% DMSO in saline (250 mg in 5 mL DMSO plus 120 mL saline), sonicating the suspension and storing it in single-use aliquots. Each mouse received a fixed injection volume of 100 µL, made up of the stock volume required for its body weight at 3 µg/g and saline to volume; across the body weights recorded during the injection period (23.2–48.4 g) this corresponds to 35–73 µL of stock and a DMSO content of 1.4–2.9% (v/v) in the injected volume.

### 2.5. Isocaloric pair-feeding

Restriction was implemented as a daily yoked ration rather than as a fixed percentage of *ad libitum* intake. Each PF/PF mouse was paired with an LF mouse of matching body-weight rank, and on each day received a weight of high-fat diet calculated as the previous day’s intake of its partner scaled by the ratio of the two energy densities (approximately 0.8), so that the pair was matched on energy rather than on mass. When the HF/PF group entered restriction at week 5 (25 August 2023), each of its mice was assigned the ration of one PF/PF mouse, and therefore inherited the same LF partner. Daily intake was taken as the difference between the weight offered and the weight remaining; spillage was not recovered separately.

### 2.6. CCK-induced satiation test

Responsiveness to cholecystokinin was tested during weeks 11 and 12, between 5 and 12 October 2023. Mice were fasted for 6 h, injected intraperitoneally with either saline or sulfated CCK-8 (3 µg/kg body weight; Bachem, Bubendorf, Switzerland) and given pre-weighed food immediately afterwards; intake was recorded at 20, 40 and 60 min and converted to energy. Every mouse was tested under both conditions in a crossover with at least 48 h between them. Because each animal served as its own control, the quantity carried into the analysis was the difference between its two intakes, saline minus CCK-8, which removes the between-animal variation in baseline intake that otherwise dominates this assay; these differences were compared across groups by one-way ANOVA separately at each time point. One mouse per incomplete pair was dropped, leaving 31 animals (LF 7, HF 8, HF/PF 8, PF/PF 8).

### 2.7. Sample collection

Mice were killed on 14 and 15 October 2023. Food was withdrawn at about 07:00 and procedures began at 13:00, giving a fast of approximately 6 h, with the order of animals arranged so that groups were interleaved rather than processed in blocks. Tail-vein blood was sampled for glucose by glucometer before euthanasia by CO2 inhalation. Blood was then drawn by cardiac puncture and plasma separated at 1,000 × g for 10 min at 4 °C. Liver, epididymal and retroperitoneal fat depots and cecum were dissected and weighed; adipose and ileal tissue were frozen on dry ice and held at −80 °C. The retroperitoneal depot of one LF mouse could not be recovered intact and is missing from the visceral fat calculation.

### 2.8. Plasma analyses

Total cholesterol, triglycerides, alanine aminotransferase and aspartate aminotransferase were determined on a Hitachi 7180 analyzer (Hitachi, Tokyo, Japan) by Dooyeol Biotech (Seoul, Republic of Korea). Two of the plasma samples were visibly hemolyzed. Hemolysis releases both transaminases from erythrocytes and so raises their measured activity without any hepatocellular injury, and the affected samples did not fall evenly across the four groups; we therefore place ALT and AST in the supplementary material and draw no inference from them. Cholesterol and triglycerides are unaffected by hemolysis at this level and are reported for every sample obtained.

### 2.9. Adipose histology

Epididymal fat was immersion-fixed in 10% neutral buffered formalin for 24 h, processed to paraffin, cut at 5 µm and stained with hematoxylin and eosin. Two fields were captured per animal at 10× on a Ki3000 microscope (Korea Lab Tech, Seongnam, Republic of Korea) and ten adipocyte diameters were traced in each with ImageJ (NIH, Bethesda, MD, USA), so that each of the 32 mice contributes 20 measurements and the mean of those is the value plotted.

Selection of the micrographs reproduced in Fig. 3A was made on the measurements rather than by eye: for each group we took the animal whose mean diameter was nearest the group mean, and within that animal the field carrying no measurement annotation.

### 2.10. Quantitative real-time PCR

RNA was recovered from epididymal fat, ileum, duodenum and liver using TRIzol (Invitrogen, Carlsbad, CA, USA) and reverse-transcribed with TOPscript RT DryMIX (dT18 plus; Enzynomics, Daejeon, Republic of Korea). Reactions used TOPreal qPCR 2× PreMIX SYBR Green (Enzynomics) on a StepOnePlus instrument (Thermo Fisher Scientific, Waltham, MA, USA); primer sequences appear in Supplementary Table S2. *Gapdh* served as the reference and expression was expressed as 2^(−ΔΔCt) against the LF group mean [15].

In line with published reporting standards for quantitative PCR [16], all statistics were computed from the raw Ct values rather than from quantities reported by the instrument software. Assays were excluded from the figures where the mean Ct exceeded 33, indicating amplification too late to quantify reliably, or where the slope of target Ct regressed on *Gapdh* Ct fell below 0.70. The second criterion tests whether target and reference share their technical variance: a slope near 1 means the two move together, so that subtracting the reference removes the shared component, whereas a slope near 0 means the reference varies independently and normalization introduces variation that is not biological. The 0.70 value is not an established community standard; it was chosen as a conservative division between these two regimes and applied uniformly to every assay, and Supplementary Table S1 reports the slope, its coefficient of determination and the mean Ct for each one so that a different threshold can be applied by the reader. Amplification efficiency was not determined for each assay, and the 2^(−ΔΔCt) method assumes that target and reference amplify with comparable efficiency. A single reference gene was used and no reference-gene stability panel was run, so instability of the reference could be detected but not corrected. Both criteria were applied identically in every tissue; no tissue-level rule was used. *Gapdh* Ct itself differed between groups in adipose tissue (P = .001) and liver (P = .020), but not in ileum (P = .09) or duodenum (P = .12); with a single reference gene we cannot tell whether that variation is technical or biological, so ΔCt results are reported as exploratory wherever the reference varied. Within each tissue the included assays were tested separately by one-way ANOVA on ΔCt and the resulting P values were adjusted across the panel by the Benjamini–Hochberg procedure [25]; adjusted values are given with the results. Exclusion and interpretation are separate decisions here: three of the eleven hepatic assays met the criteria above, but the hepatic panel is not interpreted anywhere in this report, because eight of the eleven failed them and the reference gene in that tissue varied between groups. All eleven are listed in Supplementary Table S1 for transparency.

### 2.11. Fecal microbiota analysis

Feces were collected at the end of the feeding period from four of the eight mice in each group. DNA was recovered with the ZR Fecal DNA MiniPrep kit (Zymo Research, Irvine, CA, USA) and the V3–V4 region amplified with *Bakt_341F* and *Bakt_805R* for paired-end sequencing (2 × 301 bp) on an Illumina MiSeq by Macrogen (Seoul, Republic of Korea). Reads were trimmed of adapter and primer with Cutadapt v3.2 [17] and resolved into amplicon sequence variants with DADA2 v1.18.0 [18] through filtering, denoising, merging and chimera removal, which retained between 24,862 and 53,867 reads per sample. Taxonomy came from the NCBI 16S rRNA database. Sequences were aligned with MAFFT v7.475 [19] and a tree inferred with FastTreeMP v2.1.10 [20]. Observed ASVs, Chao1, the Shannon index and Faith’s phylogenetic diversity, together with Bray–Curtis and weighted and unweighted UniFrac distances [21], were obtained with QIIME v1.9.0 [22]. Sequencing four of the eight mice in each group was decided in advance on cost grounds; the consequence for the resolution of the community comparisons is set out in Section 2.12.

### 2.12. Statistical Analyses

Values are given as mean ± SEM throughout, and the unit of analysis is the mouse. The default comparison was a one-way ANOVA across the four groups, with Tukey’s honest significant difference test used afterwards where the group effect reached significance and displayed as compact letters. Transcript data were transformed to log2 before testing, which is the same thing as testing ΔCt; bars show geometric means, and because the error bars are SEMs computed on the log2 scale and then transformed back, they are not symmetric about the bar. For body weight we also fitted a linear mixed model to every weekly measurement, with group, week and their product as fixed terms and a per-mouse random intercept, since reducing a 17-point trajectory to a single endpoint throws away both the repeated measurements and the between-animal differences in starting size.

Community composition was compared by permutational multivariate analysis of variance (PERMANOVA) [23] on Bray–Curtis dissimilarities with 9999 permutations, and the homogeneity of within-group dispersion was tested by PERMDISP [24] with the same number of permutations; a significant PERMANOVA is interpretable as a shift in community centroid only when PERMDISP is not significant. Pairwise PERMANOVA was performed between all six pairs of groups and P values were adjusted by the Benjamini–Hochberg procedure [25]. With four mice per group there are 35 distinct partitions of eight samples into two groups of four, so the smallest attainable permutation P value in a pairwise test is 1/35 = 0.029 and no pairwise contrast can reach significance after adjustment for six comparisons; pairwise results are therefore reported with this limit stated.

Analyses were performed in Python 3 with SciPy and statsmodels; PERMANOVA, PERMDISP and principal coordinates analysis were implemented directly from the distance matrices. Statistical significance was set at P < .05.

## 3. Results

### 3.1. High-fat hyperphagia was modest and pair-feeding matched intake to controls

All 32 mice survived to the end of the study. Body weight at the start of treatment did not differ between groups (26.8–27.0 g, P = 1.00; Fig. 2A).

**Figure 2.**
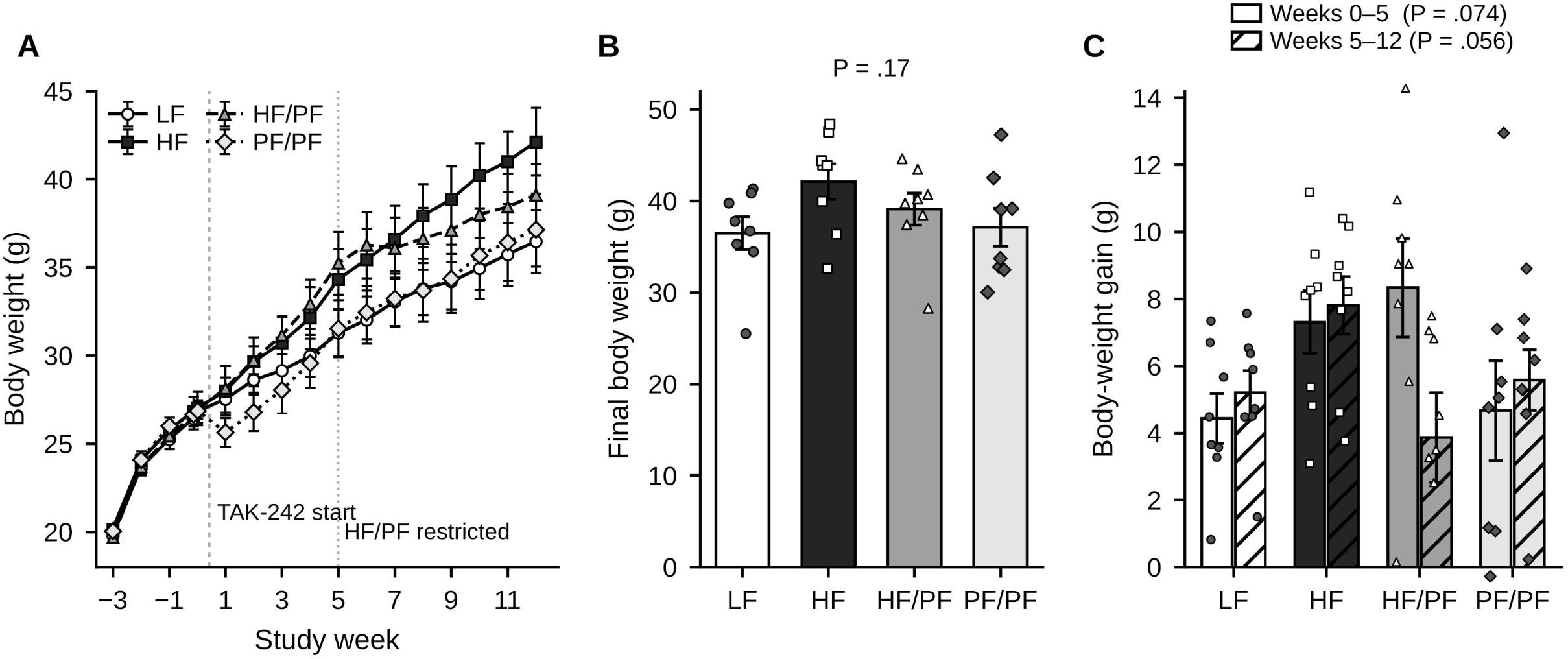
Body weight. (A) Body weight at each weighing. Symbols are group means ± SEM; the dashed line marks the start of TAK-242 injections and the dotted line the start of restriction in HF/PF at week 5. Week 0 is the day of group assignment; weights recorded before assignment are plotted at negative week numbers. (B) Body weight at the final weighing (week 13). (C) Weight gained over weeks 0–5 (open bars) and over weeks 5–12 (hatched bars). Bars are mean ± SEM with individual mice overlaid; P values in (B) and (C) are one-way ANOVA. The group × week interaction reported in the text for panel (A) is from a linear mixed model with mouse as a random intercept.

During the three acclimation weeks, when every mouse ate the low-fat diet, energy intake did not differ between the groups that were later assigned to different diets (LF 98.3 ± 2.5, HF 98.8 ± 2.5, HF/PF 96.6 ± 2.7, PF/PF 96.7 ± 1.5 kcal/week; P = .88; Fig. 1B, C), confirming that allocation did not bias intake. Over the 12 weeks on the assigned diets the *ad libitum* high-fat group ate 5.5% more energy than the low-fat control, and the groups did not differ significantly (LF 96.6 ± 3.1, HF 101.9 ± 2.9, HF/PF 98.0 ± 2.6, PF/PF 96.1 ± 3.0 kcal/week; P = .50).

Hyperphagia on the high-fat diet was therefore modest in this cohort; the excess over the low-fat control was largest between weeks 7 and 10, at 9–12%, and smaller before and after (Fig. 1B). Both pair-fed groups tracked the low-fat control throughout, as the yoked design intended.

### 3.2. Body-weight trajectories differed despite nonsignificant differences in final weight

Final body weight did not differ between groups (LF 36.5 ± 1.8, HF 42.1 ± 1.9, HF/PF 39.1 ± 1.8, PF/PF 37.1 ± 2.1 g; P = .17; Fig. 2B), and neither did weight gained over weeks 0–5 (P = .074) or weeks 5–12 (P = .056; Fig. 2C).

A linear mixed model fitted to the full trajectory resolved a difference that the endpoint comparison did not: relative to LF, the HF group gained 0.480 g/week more (P = .012), whereas neither pair-fed group differed from LF (HF/PF +0.193 g/week, P = .31; PF/PF +0.171 g/week, P = .37; Fig. 2A).

### 3.3. Adiposity measures did not differ significantly, whereas cecum mass varied among groups

Mean adipocyte diameter did not differ significantly between groups (LF 68.2 ± 3.8, HF 78.5 ± 2.5, HF/PF 77.4 ± 2.3, PF/PF 70.8 ± 3.3 µm; P = .063; Fig. 3A, B), nor did visceral fat mass expressed as a percentage of body weight (5.24 ± 0.43 to 6.81 ± 0.62%; P = .15; Fig. 3C). In both measures the two pair-fed groups lay between LF and HF without being separable from either.

**Figure 3.**
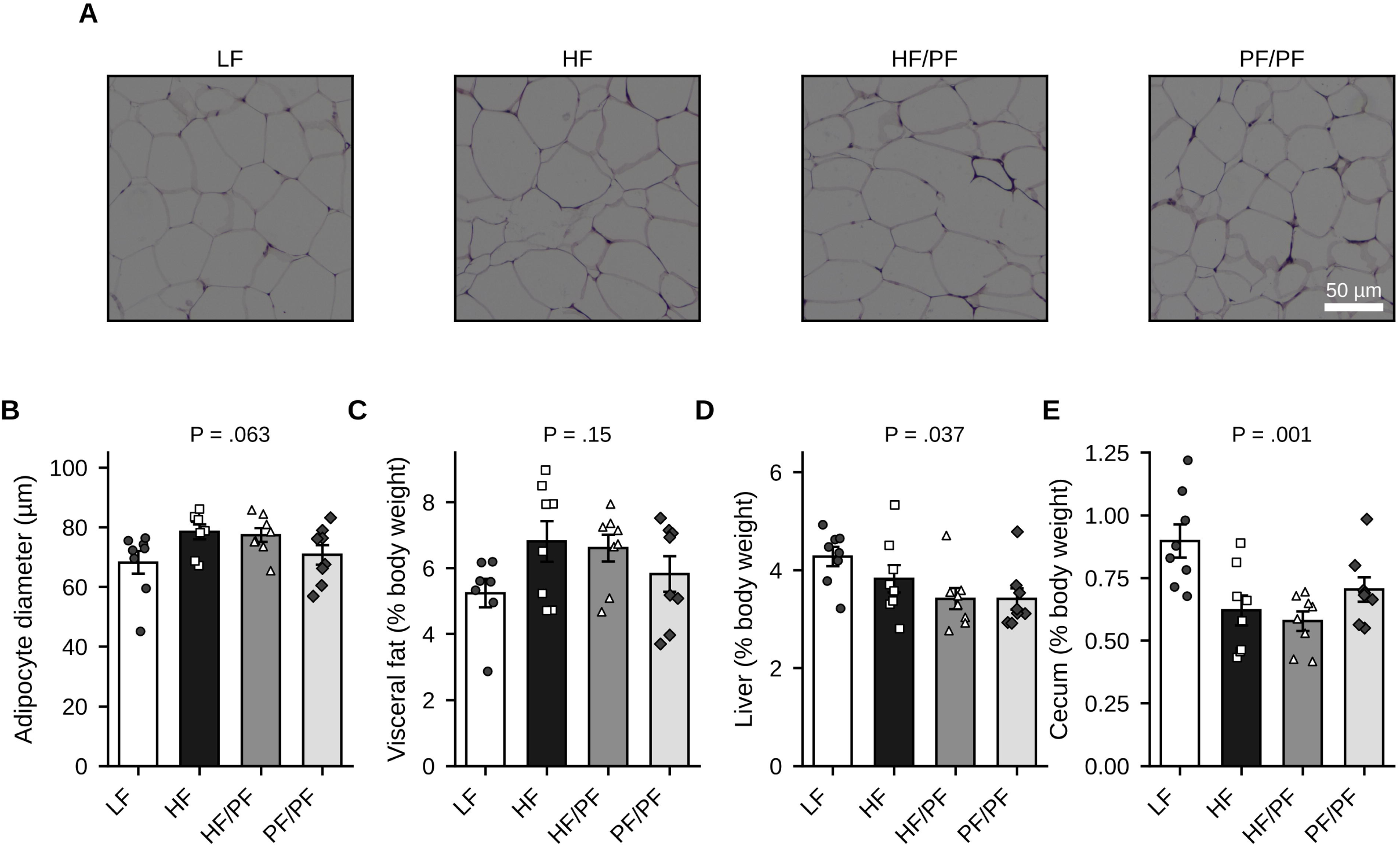
Adipose tissue morphology and organ mass. (A) Representative hematoxylin–eosin sections of epididymal adipose tissue. For each group, the mouse whose mean adipocyte diameter lay closest to the group mean was chosen (LF5, HF1, HF/PF4, PF/PF5; within 0.3–3.1 µm of the respective group mean), and within that mouse an annotation-free field of the archived micrographs. All images were acquired at the same magnification. Scale bar, 50 µm. (B) Mean adipocyte diameter, measured in ten cells per field in two fields per mouse for all 32 mice. (C) Combined retroperitoneal and epididymal fat mass,(D) liver mass and (E) cecum mass, each as a percentage of final body weight. Bars are mean ± SEM with individual mice overlaid; P values are one-way ANOVA. n = 8 per group except (C), where retroperitoneal fat was not recovered from one LF mouse (n = 7).

Cecum mass differed between groups (P = .0015; Fig. 3E) and was largest in the low-fat control (0.897 ± 0.067% of body weight) and smallest in the two groups that remained on high-fat diet (HF 0.621 ± 0.060, HF/PF 0.578 ± 0.039%; LF vs HF P = .007, LF vs HF/PF P = .002); PF/PF was intermediate (0.705 ± 0.049%) and not separable from LF (P = .083). Liver mass as a percentage of body weight varied between groups by one-way ANOVA (P = .037) but no pairwise contrast reached significance (smallest Tukey P = .055).

### 3.4. Pair-fed mice had lower fasting glucose than *ad libitum* high-fat-fed mice

Fasting blood glucose differed between groups (P = .013; Fig. 4A). Both pair-fed groups were lower than the *ad libitum* high-fat group (HF 187.9 ± 8.2 vs HF/PF 147.5 ± 9.7 mg/dL, P = .029; vs PF/PF 144.9 ± 11.1 mg/dL, P = .018), but the high-fat group did not differ from the low-fat control (163.6 ± 9.8 mg/dL, P = .29).

**Figure 4.**
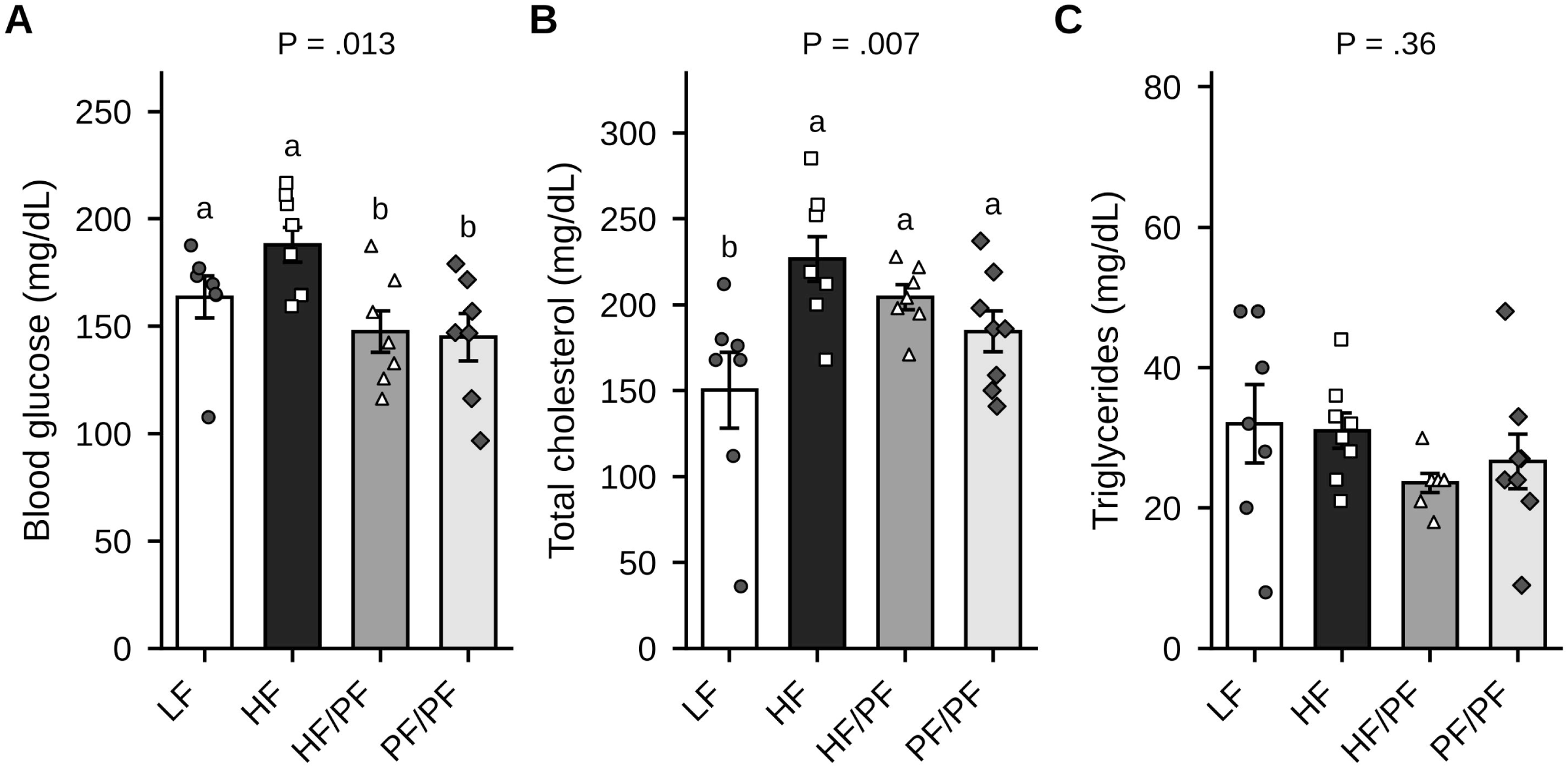
Fasting glucose and plasma lipids. (A) Fasting blood glucose, (B) total cholesterol and (C) triglycerides at sacrifice. Bars are mean ± SEM with individual mice overlaid; P values are one-way ANOVA. Where the group effect was significant, groups that do not share a letter differ at P < .05 by Tukey’s honest significant difference test; no letters are shown where the group effect was not significant. n = 7–8 per group; plasma was not available for every mouse (glucose: LF 7, HF 8, HF/PF 7, PF/PF 7; cholesterol and triglycerides: LF 7, HF 8, HF/PF 7, PF/PF 8).

Total cholesterol also differed between groups (P = .0069; Fig. 4B). The only significant contrast was HF versus LF (226.6 ± 13.0 vs 150.3 ± 22.1 mg/dL, P = .005); HF/PF (204.4 ± 7.2 mg/dL) and PF/PF (184.5 ± 11.9 mg/dL) were not separable from either LF (P = .071 and P = .35) or HF (P = .70 and P = .17). Triglycerides did not differ between groups (P = .36; Fig. 4C).

### 3.5. CCK-induced feeding suppression did not differ among dietary groups

CCK-8 reduced short-term energy intake in every group (Fig. 5A–C). The magnitude of suppression, computed within each mouse as the difference between its saline and CCK intake, did not differ between groups at any time point (20 min P = .23, 40 min P = .24, 60 min P = .58; n = 31). At 20 min, suppression was 0.65 ± 0.17 (LF), 0.82 ± 0.31 (HF), 0.42 ± 0.56 (HF/PF) and 1.56 ± 0.42 kcal (PF/PF; Fig. 5D). The apparent loss of responsiveness in the HF/PF group reflects a single animal in which intake after CCK exceeded intake after saline by 1.26 kcal; the group mean remains positive and its confidence interval overlaps every other group.

**Figure 5.**
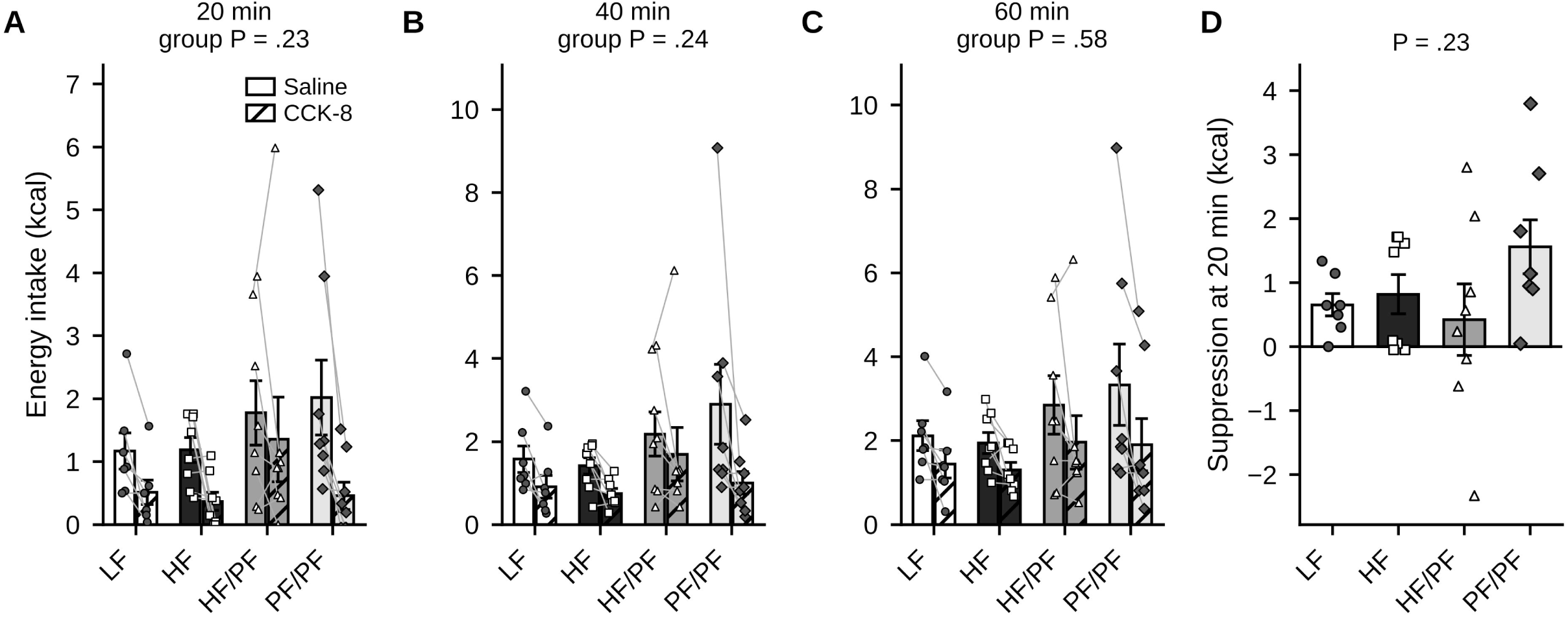
Cholecystokinin-induced satiation. Each mouse received intraperitoneal saline and, one week later, CCK-8, and its intake was recorded after both injections (crossover design). (A–C) Cumulative energy intake 20, 40 and 60 min after injection. Open bars, saline; hatched bars, CCK-8. Bars are group means ± SEM; thin lines join the two values from the same mouse. P values test the group effect on CCK- induced suppression by one-way ANOVA. (D) Suppression at 20 min, computed within each mouse as intake after saline minus intake after CCK-8; a positive value denotes suppression. Bars are mean ± SEM with individual mice overlaid; P value is one-way ANOVA. n = 31 (LF 7, HF 8, HF/PF 8, PF/PF 8); mice for which only one of the two injections was recorded are excluded.

### 3.6. Adipose *Ccl2* expression differed among dietary groups

Of the assays that met the quality-control criteria, only *Ccl2* in epididymal adipose tissue differed between groups (fold change relative to LF: HF 4.94, HF/PF 2.81, PF/PF 1.73; P = .043; Fig. 6A). *Tlr4* and *Myd88* showed the same ordering without reaching significance (P = .052 and P = .079), and *Lep*, *Srebf1*, *Pparg*, *Lcn2* and *Il1b* did not differ (P = .12–0.82). In the ileum, none of *Muc2*, *Ocln*, *Tjp1* or *Myd88* differed between groups (P = .053–0.74; Fig. 6B). In pairwise comparison only the *ad libitum* high-fat group differed from the low-fat control (4.9-fold, Tukey P = .033); no other contrast reached significance. Nine adipose assays passed quality control and were tested separately, and no gene survived Benjamini–Hochberg adjustment across that panel (smallest q = .23), so the *Ccl2* result is reported as exploratory.

**Figure 6.**
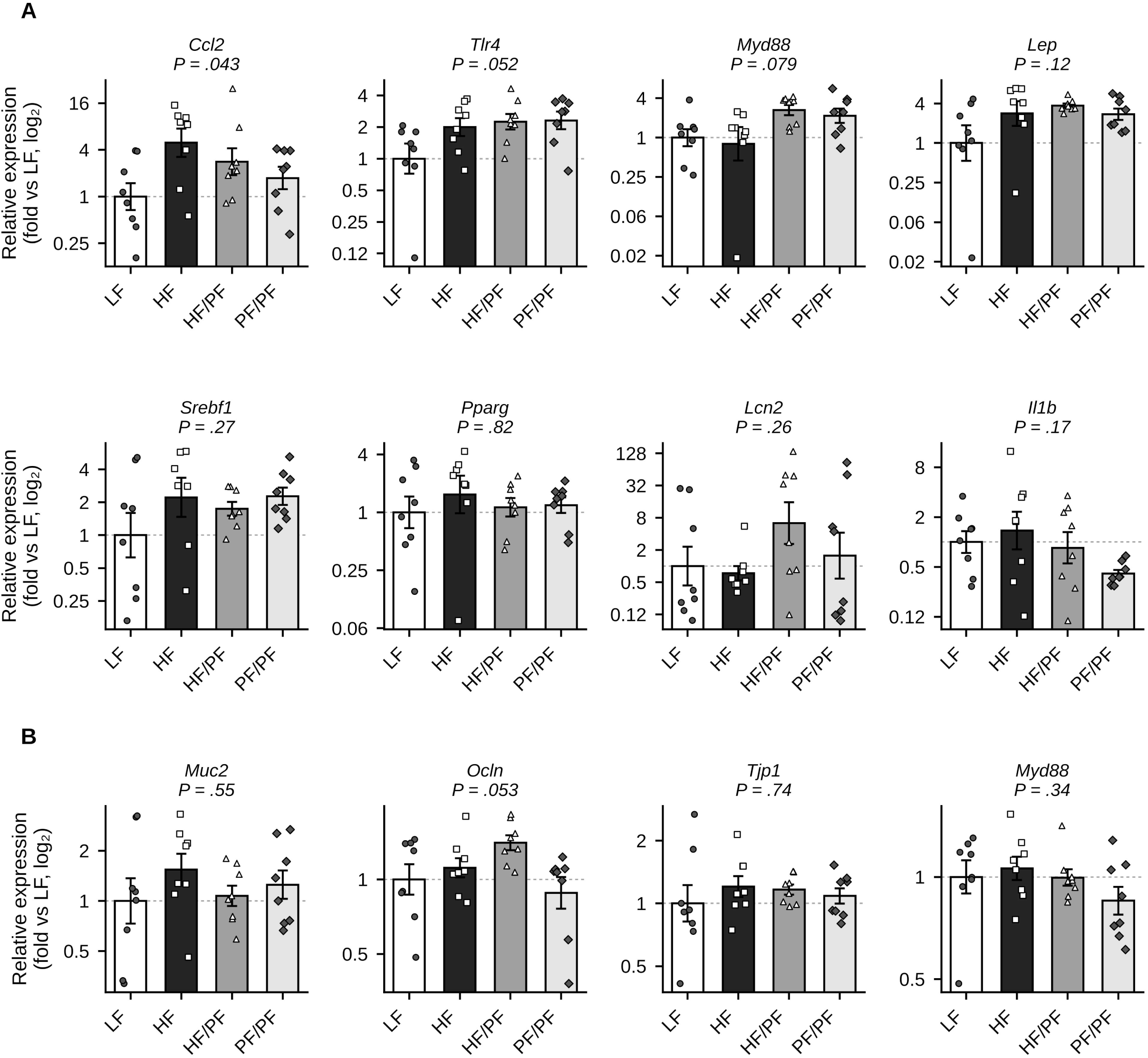
Relative gene expression in epididymal adipose tissue and ileum. (A) Adipose tissue and (B) ileum. Expression is given relative to *Gapdh* and scaled to the low-fat group mean (2^(−ΔΔCt)). Bars are the geometric mean of the group; error bars are the SEM of log₂ values back-transformed, so they are asymmetric on the linear scale. Individual mice are overlaid and the y axis is log₂ with a reference line at 1. P values are one-way ANOVA on log₂ values, equivalent to analysis of ΔCt. Gene symbols follow MGI nomenclature; *Ccl2* and *Srebf1* are referred to as MCP-1 and SREBP-1c elsewhere in the literature. Assays that did not meet the quality-control criteria given in Section 2.10 are not shown; Supplementary Table S1 lists every assay run, with its mean Ct, the slope of target Ct on *Gapdh* Ct, and the resulting verdict. n = 8 per group except where an individual reaction failed to amplify (Supplementary Table S1).

Assays were excluded where the mean Ct exceeded 33 or where the slope of target Ct regressed on *Gapdh* Ct fell below 0.70, applied identically in every tissue; Supplementary Table S1 lists all 45 assays run across the four tissues with their quality-control statistics. Nine of twelve adipose and six of eighteen ileal assays met both criteria. In liver only three of eleven did, and *Gapdh* there also differed between groups (21.9, 22.0, 22.4 and 24.4 in LF, HF, HF/PF and PF/PF; P = .020); two surviving assays showed nominal differences (*Myd88* P < .001, *Pparg* P = .012), but with most of the panel failing and the reference shifting we report no hepatic result. *Gapdh* also varied in adipose tissue (P = .001), so the adipose results below are exploratory.

### 3.7. Gut microbial community composition differed despite comparable alpha diversity

Alpha diversity did not differ between groups by any index (observed ASVs P = .18, Shannon P = .34, Faith’s phylogenetic diversity P = .19; Fig. 7A).

**Figure 7.**
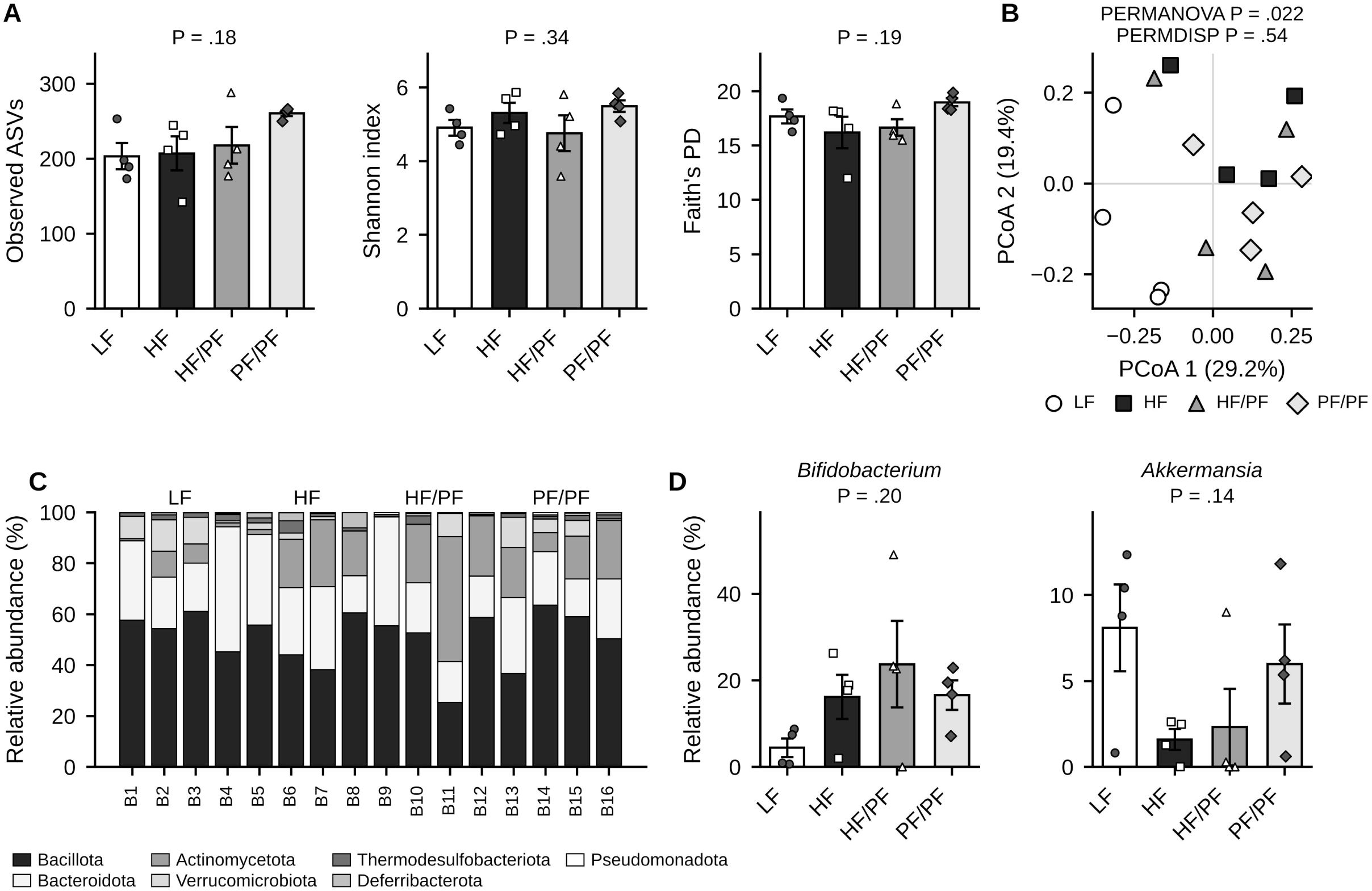
Fecal microbiota. Fecal samples from four mice per group were sequenced (16S rRNA gene, V3–V4; 24,862– 53,867 non-chimeric reads per sample). (A) Alpha diversity: observed ASVs, Shannon index and Faith’s phylogenetic diversity. Bars are mean ± SEM with individual mice overlaid; P values are one-way ANOVA. (B) Principal coordinates analysis of Bray–Curtis dissimilarities; axis labels give the variance explained by each coordinate. PERMANOVA tests for a difference in community composition between groups and PERMDISP for homogeneity of within-group dispersion, each with 9999 permutations. (C) Phylum-level relative abundance of each mouse, grouped by diet. (D) Relative abundance of *Bifidobacterium* and *Akkermansia*. Bars are mean ± SEM with individual mice overlaid; P values are one-way ANOVA. n = 4 per group throughout.

Community composition differed among groups. PERMANOVA on Bray–Curtis dissimilarities gave pseudo-F = 1.77, P = .022 (Fig. 7B), and the dispersion of each group about its centroid was homogeneous (PERMDISP F = 0.74, P = .54), indicating a shift in community centroid rather than a difference in within-group variability. In pairwise comparisons the low-fat control was the only group to differ from the others before correction (P = .029 against each of the three high-fat-origin groups), but no pairwise contrast remained significant after correction for multiple testing (q = .058); contrasts among the three high-fat-origin groups were far from significance (P = .45–.66, q = .66). As set out in Section 2.12, the smallest attainable permutation P value at this sample size is .029, so every pairwise test lay at the floor of its range and the global test is the only one that can be interpreted. The absence of a detectable difference among the three high-fat-origin groups at four mice per group is not evidence that their communities are equivalent.

At genus level, *Bifidobacterium* and *Akkermansia* varied several-fold between groups without reaching significance (P = .20 and P = .14; Fig. 7D).

## 4. Discussion

We hypothesized that, during continuous TAK-242 administration, matching the energy intake of high-fat-fed mice to that of low-fat-fed controls would normalize adiposity, intestinal barrier gene expression and the fecal microbial community. The findings supported that hypothesis only in part. Hyperphagia on the high-fat diet was modest in this cohort and pair-feeding held both restricted groups at the intake of the low-fat control; over the full trajectory the *ad libitum* high-fat group gained 0.48 g/week more than controls while neither pair-fed group differed from them, and both pair-fed groups ended with lower fasting glucose than the *ad libitum* high-fat group. Adipocyte diameter, visceral fat proportion, ileal barrier gene expression, the satiating response to CCK and microbial alpha diversity did not differ significantly among groups. Among the transcript results reported, adipose *Ccl2* was the only gene showing a nominally significant group difference, which did not survive correction for multiple testing; of all community measures, only composition differed. These observations fit a model in which energy intake and diet composition act on different components of the high-fat-diet phenotype better than one in which the phenotype responds as a whole to energy availability.

The contrast on which the design depends was smaller than intended, for a reason that lies in the cohort rather than in the feeding protocol. The *ad libitum* high-fat group ate 5.5% more energy than the low-fat control over the 12 weeks on the assigned diets, and the group difference was not significant; the excess reached 9–12% only between weeks 7 and 10. Pair-feeding itself worked as designed, holding both restricted groups at the intake of the low-fat control from the week each entered restriction, and intake did not differ between the groups during the three acclimation weeks, when all mice ate the low-fat diet. What was small, in other words, was the quantity being matched away, and because feeding pattern itself shapes microbial composition, inflammation and gut-brain signaling in high-fat-fed rodents [26], that magnitude bears on every outcome below. In our preceding experiment the *ad libitum* high-fat group ate substantially more than the low-fat control [3], and the weaker weight phenotype reported below is the most likely consequence of the narrower intake difference here. Because TAK-242 was given to every animal from week 0.4, any effect of the compound on feeding applies equally to all four groups and cannot be separated from diet; intake in the low-fat group was stable from acclimation to the end of the study, which argues against a large anorectic effect, but does not exclude one.

Body weight differed in trajectory but not at the endpoint. Final weight did not differ, and neither did weight gained in either half of the study; only the mixed-model analysis of the full trajectory resolved a difference. Endpoint and trajectory disagree because a single final weight discards the repeated measurements and absorbs the large between-animal variation in starting size, which the random intercept removes. We report both and claim no effect of diet on final body weight. The weight phenotype is weaker than in our preceding experiment [3], in which restriction normalized a clear weight gain under *ad libitum* high-fat feeding, and the reduced intake contrast described above is the most likely reason. Fasting glucose was lower in both pair-fed groups than in the *ad libitum* high-fat group, but the low-fat control itself sat high enough that it did not differ from the *ad libitum* high-fat group, so this is a comparison among high-fat-fed groups rather than evidence that high-fat feeding raised glucose.

The adipose measures ordered the groups as expected without reaching significance. Adipocyte diameter and visceral fat proportion were largest in the two groups of high-fat *ad libitum* origin and smallest in the low-fat control (P = .063 and P = .15); with eight mice per group the study was not powered to resolve differences of this size, so these results are consistent with, rather than evidence for, the dissociation between body weight and adiposity that we reported previously [3], which parallels normal-weight obesity in humans [27]. Adipose *Ccl2*, the only transcript to differ among groups before adjustment for the nine assays tested, followed the same ordering: highest in the *ad libitum* high-fat group, intermediate in the group pair-fed from week 5, and close to the low-fat control in the group pair-fed from week 0. This parallels the persistence of adipose *Tnf* expression we observed in the treatment model previously [3] and is consistent with evidence that obesity-associated immune and transcriptional features of adipose tissue persist after weight normalization [4–6]. The difference rested on the contrast between the *ad libitum* high-fat group and the low-fat control and did not survive adjustment across the panel, so we treat it as exploratory. CCL2 recruits macrophages to adipose tissue and contributes to insulin resistance and hepatic steatosis in obese mice [28], and we have found in the same model that adipocyte hypertrophy, the fecal community and cholecystokinin responsiveness can shift together under a dietary intervention [29]. *Tlr4* and *Myd88* ordered identically without reaching significance; during TAK-242 administration their transcript abundance is in any case difficult to interpret, since the compound acts on signaling rather than on transcription.

Neither intestinal barrier gene expression nor CCK-induced satiation differed significantly among groups. None of *Muc2*, *Ocln*, *Tjp1* or *Myd88* differed in the ileum, and the suppression of short-term intake by CCK-8 was similar across groups at every time point. Both outcomes replicate our earlier findings in the same model without the inhibitor [3], and the barrier result is consistent with reports that diet-induced metabolic impairment can develop without loss of barrier integrity [30]. Ileal barrier gene expression in high-fat-fed rodents is nonetheless responsive to dietary intervention in our hands [31,32], so the absence of a detectable change here is unlikely to reflect an insensitive readout. The CCK result also corrects an earlier internal analysis of this experiment, which compared saline and CCK within each group separately and concluded that the treatment group had lost responsiveness; in a crossover design the quantity that answers the question is the within-animal difference between injections, on which the groups did not differ. The apparent loss reflected a single animal that ate more after CCK than after saline.

The pattern of between-group variation in the fecal community was more closely associated with diet composition than with energy intake. Alpha diversity did not differ by any index, but community composition did, and homogeneous within-group dispersion indicated a shift in the position of the community centroid rather than a difference in variability. Before correction for multiple testing the low-fat control was the only group that differed from the others; no pairwise contrast survived correction, and contrasts among the three high-fat-origin groups were far from significance. At four mice per group the pairwise tests lay at the floor of the permutation distribution, so the absence of a detectable difference among those groups is not evidence that their communities are equivalent, and the global test is the only one that can be interpreted. What can be said is that pair-feeding, whether begun at week 0 or at week 5, did not move the community toward the low-fat profile, which accords with reports that a high-fat diet is associated with community composition independently of obesity [33–35] and that energy restriction reshapes the microbiota in a direction of its own [36,37]. Cecum mass followed the same ordering and offers a mechanistic reading: largest in the low-fat control, smallest in the two groups still eating the high-fat diet, and intermediate in the group pair-fed throughout. Cecal enlargement reflects the quantity of fermentable substrate reaching the hindgut, and the low-fat diet supplies several times more corn starch than the high-fat diet, so the relevant dietary variable may be fermentable carbohydrate rather than fat as such — consistent with reports that many microbial effects attributed to high-fat diets are reproduced by removing fermentable fiber [38,39], with our own observations on fermentable substrates added to a high-fat diet [40,41], and with the role of resistant starch as hindgut substrate [42]. Energy restriction reduces the absolute amount of every dietary component but leaves their proportions unchanged, which is what a composition-driven effect would require. *Bifidobacterium* and *Akkermansia* varied several-fold without reaching significance [43].

Read alongside our preceding experiment [3], the two studies agree on where the boundary between energy intake and diet composition falls. In both, the measures that followed energy intake were body weight and circulating markers, and the measures that remained associated with diet composition were the proportion of body weight stored as visceral fat, an adipose inflammatory transcript, and the structure of the fecal community; in both, CCK-induced satiation and ileal barrier gene expression were unaffected by either variable. That the same boundary appears in two independently run cohorts is the strongest statement these data support, and it is stronger than either experiment makes alone. The comparison is nonetheless an observation across experiments rather than a controlled contrast, since the two differ in animal batch, season, diet lot and in more than thirty intraperitoneal injections as well as in the inhibitor. What the present experiment adds is that the boundary is not abolished when TAK-242 is given to every animal throughout: the features that resisted energy restriction without the compound also resisted it during its administration.

Several limitations should be noted. The absence of a vehicle-injected arm is the principal one: no comparison here isolates the effect of TLR4 inhibition, and target engagement was not verified, so the degree and persistence of inhibition over 12 weeks are unknown. Restriction of the group pair-fed throughout began two days before the first injections, so restriction and the start of dosing cannot be separated. Hyperphagia on the high-fat diet was modest, which narrowed the contrast the design depends on. With eight mice per group three outcomes fell between P = .05 and P = .08, and absence of significance is not evidence of absence. The microbiota was analyzed in four mice per group at one time point, at which pairwise contrasts cannot reach significance. The reference gene is the weakest part of the transcript analysis. *Gapdh* Ct differed between groups in both adipose tissue (P = .001) and liver (P = .020), and with a single reference gene and no assay-level efficiency determination we cannot tell whether that variation is technical or biological. The same criteria were applied in every tissue; liver is not interpreted because eight of its eleven assays failed them, not because a different rule was used. The adipose results, *Ccl2* included, are exploratory for the same reason, and the slope criterion is a pragmatic diagnostic rather than an established standard. Plasma transaminases are reported without interpretation because two samples were hemolyzed. Finally, only male mice were studied, and body composition, energy expenditure and insulin sensitivity were not measured. Further studies are warranted in which the inhibitor, and not only the diet, is varied between groups, target engagement is confirmed, and fermentable substrate is manipulated independently of dietary fat.

## 5. Conclusion

During continuous TAK-242 administration, matching the energy intake of high-fat-fed mice to that of low-fat-fed controls prevented the steeper weight trajectory of *ad libitum* high-fat feeding and was accompanied by lower fasting glucose, even though hyperphagia on the high-fat diet was modest in this cohort, but adipocyte diameter, visceral fat proportion, ileal barrier gene expression, CCK-induced satiation and microbial alpha diversity did not differ significantly among groups, and the fecal community did not shift toward the low-fat profile. Adipose *Ccl2* expression and fecal community composition were the only measures that distinguished the groups. These findings suggest that metabolic and gut microbial responses to high-fat feeding differ in their sensitivity to energy restriction, and that dietary composition and energy intake are worth treating as distinct factors in nutritional intervention studies. Whether TLR4 signaling contributes to that separation cannot be answered by this design.

## Acknowledgment

None.

## Author Contributions

Eunji Jeong: Data curation, Formal analysis, Investigation, Methodology, Visualization, Writing – original draft. Seoyeon Chae: Data curation, Investigation, Methodology, Writing – review & editing. Sungjin Eun: Data curation, Investigation, Methodology, Writing – review & editing. Hyunwoo Chae: Data curation, Formal analysis, Investigation, Methodology, Visualization, Writing – review & editing. Sunhye Lee: Conceptualization, Formal analysis, Funding acquisition, Investigation, Methodology, Project administration, Resources, Supervision, Validation, Visualization, Writing – original draft, Writing – review & editing.

## Author Declarations

The authors declare no financial or personal conflicts of interest.

## Data availability

Raw 16S rRNA gene amplicon sequencing data generated in this study have been deposited in the NCBI Sequence Read Archive under BioProject accession number PRJNA1549769 (release date 23 October 2026); the preceding experiment is deposited under PRJNA1535893. The per-animal values underlying every figure, including untransformed Ct values and all adipocyte measurements, are provided as a source data file. All other data are available from the corresponding author on reasonable request.

## Declaration of AI and AI-assisted technologies in the writing process

During the preparation of this work, the authors used Claude (Claude Opus 5.5; Anthropic, San Francisco, CA, USA) to improve the readability and language of the manuscript. After using this tool, the authors reviewed and edited the content as needed and take full responsibility for the content of the publication.

## List of abbreviations

ANOVA: analysis of variance
ASV: amplicon sequence variant
CCK: cholecystokinin
Ct: cycle threshold
DMSO: dimethyl sulfoxide
HF: high-fat diet fed *ad libitum*
HF/PF: high-fat diet fed *ad libitum* then pair-fed
LF: low-fat diet
PCoA: principal coordinates analysis
PERMANOVA: permutational multivariate analysis of variance
PERMDISP: permutational analysis of multivariate dispersions
PF/PF: pair-fed throughout
SEM: standard error of the mean
SRA: Sequence Read Archive
TLR4: Toll-like receptor 4

